# Selection of movement rules to simulate species dispersal in a mosaic landscape model

**DOI:** 10.1101/2024.02.26.582052

**Authors:** Susannah Gold, Simon Croft, Richard Budgey, James Aegerter

**Affiliations:** National Wildlife Management Centre, Animal and Plant Health Agency, York, United Kingdom

**Keywords:** Dispersal, Movement, Spatial modelling, Kernels, Agent-based models

## Abstract

Dispersal is an ecological process central to population dynamics, representing an important driver of movement between populations and across landscapes. In spatial population models for terrestrial vertebrates, capturing plausible dispersal behaviour is of particular importance when considering the spread of disease or invasive species. The distribution of distances travelled by dispersers, or the dispersal kernel, is typically highly skewed, with most individuals remaining close to their origin but some travelling substantially further. Using mechanistic models to simulate individual dispersal behaviour, the dispersal kernel can be generated as an emergent property. Through stepwise simulation of the entire movement path, models can also account for the influence of the local environment, and contacts during the dispersal event which may spread disease. In this study, we explore a range of simple rules to emulate individual dispersal behaviour within a mosaic model generated using irregular geometry. Movement rules illustrate a limited range of behavioural assumptions and when applied across these simple synthetic landscapes generated a wide range of emergent kernels. Given the variability in dispersal distances observed within species, our results highlight the importance of considering landscape heterogeneity and individual-level variation in movement, with simpler rules approximating random walks providing less plausible emergent kernels. As a case study, we demonstrate how rule sets can be selected by comparison to an empirical kernel for a study species (red fox; *Vulpes vulpes*). These results provide a foundation for the selection of movement rules to represent dispersal in spatial agent-based models, however, we also emphasise the need to corroborate rules against the behaviour of specific species and within chosen landscapes to avoid the potential for these rules to bias predictions.

## Introduction

Abstractions of real landscapes are increasingly being integrated into ecological models to allow consideration of how spatial structure and landscape features affect processes such as disease spread (Manlove et al., 2022; White, Forester, & Craft, 2018a), range expansion of invasive species (Synes et al., 2016) and metapopulation dynamics of species of conservation concern (Sullivan, Michalska-Smith, Sperry, Moeller, & Shaw, 2021). In these models, dispersal is a key ecological process that needs to be appropriately represented to capture the movement of individuals across landscapes, and how this influences population dynamics. As with all model processes, dispersal rules should be corroborated to assure their fitness-for-purpose and to identify any potential to bias model outputs, for example where implausible movements might lead to the unrealistic spread of disease. The avoidance of this form of artefact is especially important where models are used to inform decision-making (e.g., risk assessment, policy development or management) as unrecognised bias may influence model predictions.

Dispersal here includes both natal dispersal, where maturing individuals leave their natal home range and settle to breed elsewhere, and the re-settlement of adults due to resource or reproductive competition (Bowler & Benton, 2005), but excludes daily movements within a home range. Following Bowler and Benton (2005), we divide the dispersal process into three stages, each representing decisions made by dispersers: emigration from their origin, movement, and immigration to a final settlement location. For each of these stages, numerous factors may influence individual animal behaviour, and movement itself may involve a sequence of steps, each with decision processes mediated by local factors. For example, movement behaviour might be influenced by the individual’s internal state (e.g., energy availability or health status), perceptual or navigational ability, movement ability, and biotic or abiotic properties of the environment (Nathan et al., 2008). This complexity can result in highly variable dispersal outcomes within a single species. Theoretical studies have highlighted the impact that this individual-level and environmentally determined variation can have on population-level outcomes, such as population viability and disease maintenance (Gorton & Shaw, 2022; Markov & Ivanko, 2022; Scherer et al., 2020).

Choices in modelling the dispersal process produce trade-offs across a number of principles. Fundamental amongst these include whether the purpose requires only the movement outcome to be simulated (a classic “flight” defined only by an origin, a final settlement location and the vector between them) or a mechanistic model representing the movement process itself (simulating a potentially longer walk describing the locations along the trajectory between the origin and settlement location). In mechanistic models of dispersal, the process can be simulated so that movements depend on information which would realistically be perceived by the agent, for example local habitat quality or the presence of barriers. As a result, the distance travelled by an agent during dispersal is simulated as a responsive and emergent process, rather than being pre-defined by a distribution and applied inflexibly in every case. In the case of disease models, simulation of the movement route also allows for consideration of disease transmission during the walk. Previously developed ecological models have used a range of routines to mechanistically model dispersal, from simple diffusion, assuming environmental homogeneity, to complex individual-based models allowing agents to perceive and respond to heterogenous environments (e.g., Croft, Aegerter, Beatham, Coats, & Massei, 2021; Hawkes, 2009; Kanagaraj, Wiegand, Kramer-Schadt, & Goyal, 2013; La Morgia, Malenotti, Badino, & Bona, 2011; Pauli et al., 2013; Vuilleumier & Metzger, 2006). The decision on the complexity of the simulation will depend on both the purpose of the model and the information available for parameterisation. The method of representation is of importance, as even relatively small changes to movement can influence population dynamics in models (Hawkes, 2009) and the potential for disease spread (Scherer et al., 2020; Tracey, Bevins, VandeWoude, & Crooks, 2014; White et al., 2018a) .

Predictive ecological simulations often cannot be validated in the conventional sense, through generation of new data from experimental studies, due to the complexity of the systems considered (Augusiak, Van den Brink, & Grimm, 2014; Joseph et al., 2013) or where their purpose may confound empirical science (e.g., epidemiology of notifiable disease or spread of a harmful invasive species). However, confidence in the credibility and utility of predictions can be improved by verification of the conceptual model, assuring that principal components of the model are individually free from unnecessary error and show minimal potential for bias, (e.g., Holland, Aegerter, Dytham, & Smith, 2007). For corroboration of dispersal rules in models, there is an increasing amount of animal movement data available (Neumann et al., 2015); in particular, the availability of GPS technology has led to an increase in fine-scaled tracking data, allowing consideration of the entire movement process. However, empirical studies for many species are limited to mark-recapture studies or *ad hoc* observations, which mainly inform description of dispersal flights rather than walks. Information on dispersal flights is often summarised into a dispersal kernel, a probability density function representing the Euclidian distance between the origin and final location (either the site of settlement or mortality). To validate movement rules, emergent dispersal distances from models should be compared to these empirical kernels where available. However, in the absence of suitable data, comparison to a kernel shape considered plausible can contribute to ensuring that unrealistic patterns of movement are not being generated. Typically, dispersal kernels follow a right-skewed and fat-tailed distribution, with most individuals remaining close to their origin but some long-distance dispersers settling much further away (Fandos et al., 2021; Nathan, Klein, Robledo-Arnuncio, & Revilla, 2012; Soulsbury, Iossa, Baker, White, & Harris, 2011; Whitmee, 2011). Though relatively infrequent, long-distance dispersal or long walks from the origin can play a significant role in the spread of disease or invasive species (Hastings et al., 2005; Jeltsch, Müller, Grimm, Wissel, & Brandl, 1997; Mundt, Sackett, Wallace, Cowger, & Dudley, 2009). The exact shape of kernels, and the probability of long-distance dispersal events, will be highly dependent on the species and individual landscape. Therefore, methods to simulate dispersal in spatially explicit predictions require flexibility in the approach used (Nathan et al., 2012).

In this paper, we explore a suite of simple movement rules, reflecting those commonly used in spatial models, to simulate dispersal of agents across a mosaic landscape. We use these examples to consider the emergent distance distributions, and how the choice of rules can be corroborated by comparison to realistic dispersal kernels. Our overall aim is to provide a foundation for representing dispersal for terrestrial species that can be used in spatially explicit population models, in particular for predictive modelling used by decision-makers for management of wildlife, the epizootiology of their diseases, or the establishment and spread of invasive species. As a case study, we demonstrate how movement rules could be selected for dispersal of the red fox (*Vulpes vulpes*), based on comparison to published estimates of dispersal.

## Materials and Methods

### Overview

We simulated dispersal across synthetic landscapes generated using a simple consistent method, Voronoi tessellation, to produce a mosaic of irregular parcels. A suite of parcel-to-parcel walks were run across these landscapes and the dispersal kernels they produce were studied. A sensitivity analysis was conducted to describe the flexibility of different rule sets to generate plausibly shaped kernels. We then took an example dispersal kernel based on empirical data for the red fox and demonstrated how this can be simulated within our modelling paradigm.

### Method of landscape generation

The choice of how dispersal is represented is closely tied to the method used to represent the underlying landscape. All model landscapes have statistical properties which are critical to the credibility of the model processes run across them. Many spatially explicit modelling studies use cells in regular geometries to represent landscapes and movement, primarily grid-based representations (e.g., Bocedi, Pe’er, Heikkinen, Matsinos, & Travis, 2012; Boone & Hunter, 1996; Gardner & Gustafson, 2004; La Morgia et al., 2011). However, regular geometries are limited in their ability to capture complex real-world landscapes (Holland et al., 2007). The fixed scale produces error in the representation of underlying environmental features and allows for potential bias in movement direction (Chipperfield, Holland, Dytham, Thomas, & Hovestadt, 2011).

Here we build upon earlier work which establishes that complex representations of real-world landscapes can be achieved using a tessellated mosaic of cells (hereafter parcels to emphasise that they may combine and contain varied habitats or resources) constructed using an irregular geometry (Holland et al., 2007), conceptually close to a geographic automata (Benenson & Torrens, 2003; White, Forester, & Craft, 2018b). Within this geometry, each parcel represents the geographic locations (either in real or synthetic landscapes) of sub-populations or individual agents (e.g., territories or home ranges). Parcels can be built to include accurate representation of real-world barriers or other key geographical features, and their scale can be varied within a simulated landscape where required (e.g., where territory size varies in response to resource availability). Furthermore, depending on the rules used to build parcels and the traits assigned to them (binary, categorical or continuous), they might be considered a generalised form of patch/matrix model able to represent the complex nature of dispersal to habitat specialists (Tischendorf, Bender, & Fahrig, 2003) or the experience of real-world generalists as they assemble resource from alternative and diverse habitats within most parcels across a landscape. Crucially, when using multiple instances of these randomised landscapes, the implementation of inter-parcel movement is free of directional bias (Holland et al., 2007), a feature essential to models which are designed to support decision-making that must be free from artefact (Bocedi et al., 2012). In addition, using this method, a spatially explicit process such as a dispersal walk can be run using only information about adjacent neighbours, allowing simulations to run outside costly GIS environments. The synthetic landscapes generated are essentially abstract and scale-free (i.e., consistent relationships between parcel area, inter-parcel movement and the generation of kernels), however in this paper we represent these landscapes at a set scale appropriate for considering red fox dispersal to illustrate a direct application to simulations in real-world landscapes.

For this study, virtual landscapes of 100 km by 100 km (10,000 km^2^) were divided into discrete parcels by Voronoi tessellation (See Figure 1). For simplicity in this experimental study, we assumed no barriers across a single contiguous mosaic extent. For each landscape 10,000 seed locations were distributed randomly across the arena to give a mean parcel size of 1 km^2^, and parcels were formed using the st_voronoi function (sf package in R). Due to the random nature of each landscape, dispersal walks were run in ten randomised instances of these mosaics. Statistical descriptions of these landscapes are shown in the supporting information: parcel size distribution, step length distribution, mean and range of neighbourhood metrics (mean and range of neighbours per parcel) and parcel qualities. Within this framework, we consider parcels as the space occupied by individual agents (equivalent to territories or home ranges). During dispersal, individuals move between these parcels, with the length of inter-parcel movements assumed to approximate to centroid-centroid distances. In effect, agents therefore move across a network of nodes in a Delaunay triangulation (the network linking parcel centroids).

**Figure 1:**
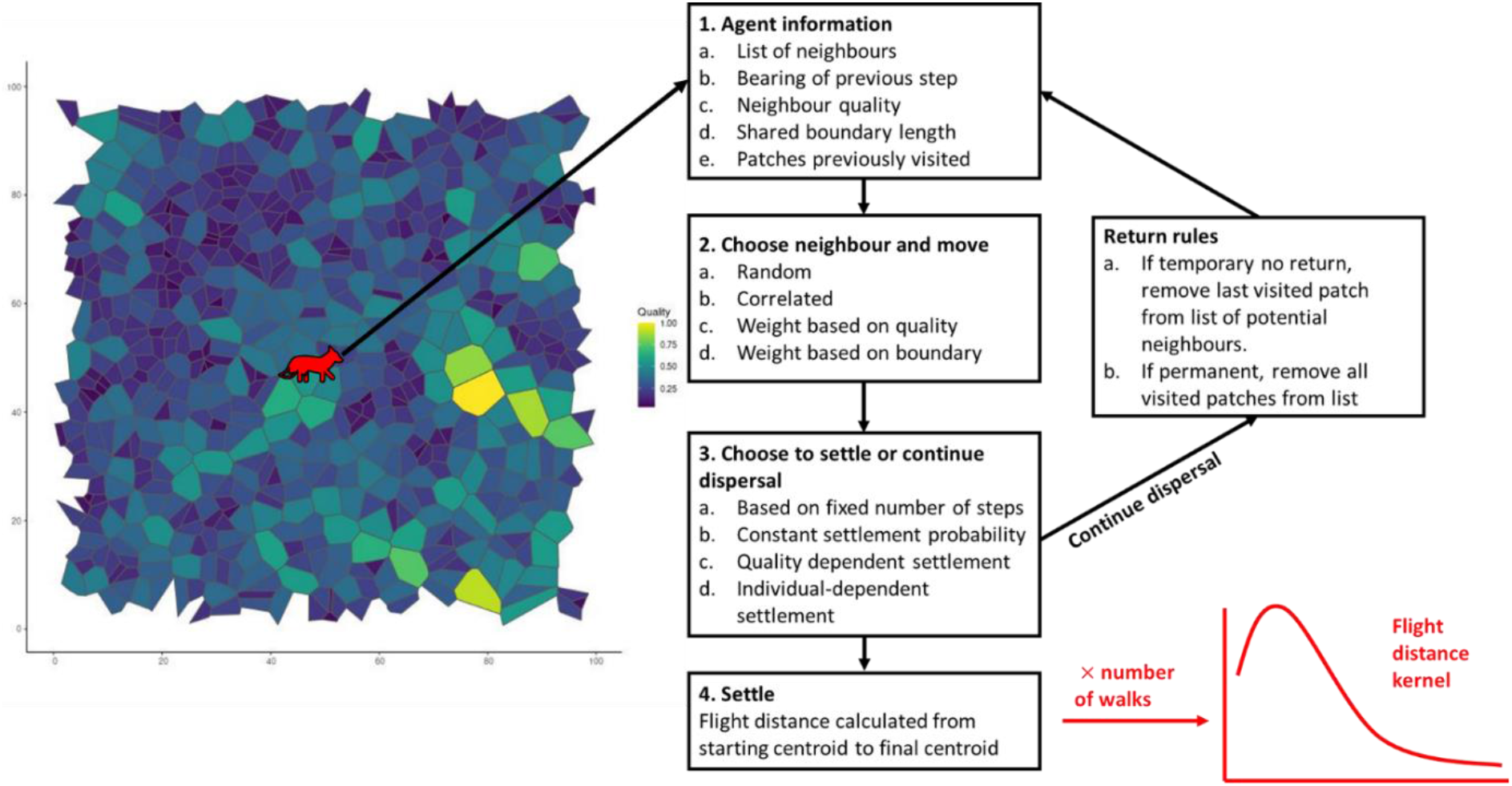
Conceptual dispersal model. A subset of a landscape extent is shown for visualisation. A full description of rules considered is shown in Table 1.

### Generation of quality patterns

Parcel characteristics may influence dispersal behaviour and consequently the emergent dispersal kernel. For example, the composition and configuration of habitats within a parcel might influence its resistance to transit (Pflüger & Balkenhol, 2014) or the likelihood of an individual choosing to end its walk and settle (Matthysen, 2005). These factors will be specific to the particular landscape. However, to demonstrate the effect heterogeneity may have on dispersal kernels, we assign each parcel within our mosaic landscapes a quality value. In this study, this represents an abstract quality that can affect both the likelihood of choosing to move into a parcel and choosing to settle within that parcel. To ensure that the spatial qualities of this variation were consistent between alternative randomisations of landscapes we generated an underlying quality description using the neutral landscape model package in R (NLMR) (Sciaini, Fritsch, Scherer, & Simpkins, 2018). This package produces raster patterns based on theoretical distributions; here we use the midpoint displacement algorithm with a default roughness of 0.5. Raster values were summed across each parcel and qualities were then normalised by dividing by the maximum value so that quality for each patch ranged between 0 and 1. In these synthetic landscapes it was assumed that all parcels were suitable for settlement, as may occur with a generalist species. The distribution of assigned parcel qualities is shown in Appendix 1 (S1): Figure S1.

**Table 1:** Dispersal movement rules simulated. Asterisks indicate the default choice when varying other rules.

| Variable | Rule | Behaviour assumption |
| --- | --- | --- |
| **1. Neighbour choice** | **Random** - Equal probability. * | No systematic behaviour. |
|  | **Correlated** - Initial step random, with each further step closest to previous bearing | Whole flight is directional, with steps informed by individual's attempt to follow a consistent direction. |
|  | **Biased - shared boundaries** - Weighted based on the length of shared boundary. | Animals likely to select most accessible neighbour (longer shared boundary). |
|  | **Biased – quality** - Weighted based on the quality. | Animals likely to select “best” neighbour based on a perceived characteristic. |
| **2. Rules on return** | **None** - Equal probability. * | Animals might immediately backtrack. |
|  | **Temporary no return** - Return to previous parcel prohibited. | Animals will not immediately backtrack. |
|  | **Persistent no return** - Any previously visited parcel is prohibited. | Constant exploration. |
| **3. Settlement probability** | **Constant length** - Move $n$ steps and settle (Default $n=10$). | All animals move through a constant number of parcels and then inevitably settle |
| | **Constant settlement probability** - Fixed and common probability of settlement at each step, $p_{\text{settlement}}$ (Default $p_{\text{settlement}}=0.1$). | All animals move and randomly settle in the current parcel based on a set behaviour. |
| | **Quality dependent settlement probability** - variable $p_{\text{settlement}}$ based on variation in parcel “quality” (parcel quality measure as a relative probability 0-1). High patch quality = high probability of settlement. Default parcel quality values in S1: Figure 1. | All animals move and settle in the current parcel dependent on its perceived quality. |
|  | **Individual settlement probability** - Each agent has a unique settlement probability, drawn from a beta distribution, that remains constant throughout the walk. As a default, a uniform distribution between 0 and 1 was used. | Inter-individual variation occurs, for example in personality or phenotypic movement ability, leading to a different tendency to settle. |

### Movement rules for simulation of dispersal kernels

Dispersal was simulated as a sequence of stepwise movements of agents between neighbouring parcels until settlement. Disperser mortality was not considered here, with only successful dispersal movements leading to settlement considered. To avoid the influence of boundaries, origins were randomly selected from central parcels (parcel centroid was ≥20 km from arena edge). Here we ignore any parcel variation, for example in size or quality, that might influence emigration and all central parcels had an equal probability of being an origin. For each step, agents choose from a list of directly adjacent neighbours, with varying probabilities depending on the rule set. As a baseline, we ran a parcel-to-parcel uncorrelated random walk for 10 steps (neighbours chosen randomly and no rules on return). Different simple rules caricaturing aspects of animal behaviours were then incorporated into movement walks (Figure 1, Table 1).

### Analysis of movement rules output

For initial exploration of the impact of movement rules, neighbour choice, return rules and settlement rules were varied independently, and the resulting kernels plotted in comparison to the default random walk. For each rule set, 1000 trials were simulated across each of 10 randomised landscapes (10,000 trials total).

To account for the influence of combining different rule sets and parameterisations, we conducted a sensitivity analysis on the influence of different settlement probability parameterisations for each rule set. All potential combinations were considered (4 neighbour choice x 3 return rules x 3 settlement probability), with the exception of the correlated walk in which, as the walk is unidirectional, return rules do not apply. Therefore, 40 combinations of rules were considered in total. For the settlement rules, parameter values were varied over a range. For the constant length walk, *n* ranged between 1-20. For the constant settlement probability p_settlement ranged from 0.1-1 (intervals of 0.05). In the case of quality dependent settlement, the kernel will be dependent on the specific landscape, and how quality varies both in magnitude and spatially. Exploration of different spatial distributions of quality was not in the scope of this paper, therefore we solely considered how variation in the relative quality influenced emergent kernels. To do this we ran the walks across a series of landscapes with the same spatial distribution of quality, but with the quality values determining settlement reduced relative to the default (shown in S1 Figure 1) using a scaling factor (0.1 to 1, with intervals of 0.05). For incorporating individual variation in settlement, probabilities were drawn from 8 different beta distributions (S1 Figure 2). These distributions vary from a uniform distribution (Distribution 1~ Beta(α=1, β=1)) where all settlement probabilities are equally likely, to distributions where either high settlement probabilities are most likely to be drawn (Distribution 5-Beta (α=5,β=1) or low probabilities (Distribution 2-Beta(α=5, β=1)).

**Figure 2:**
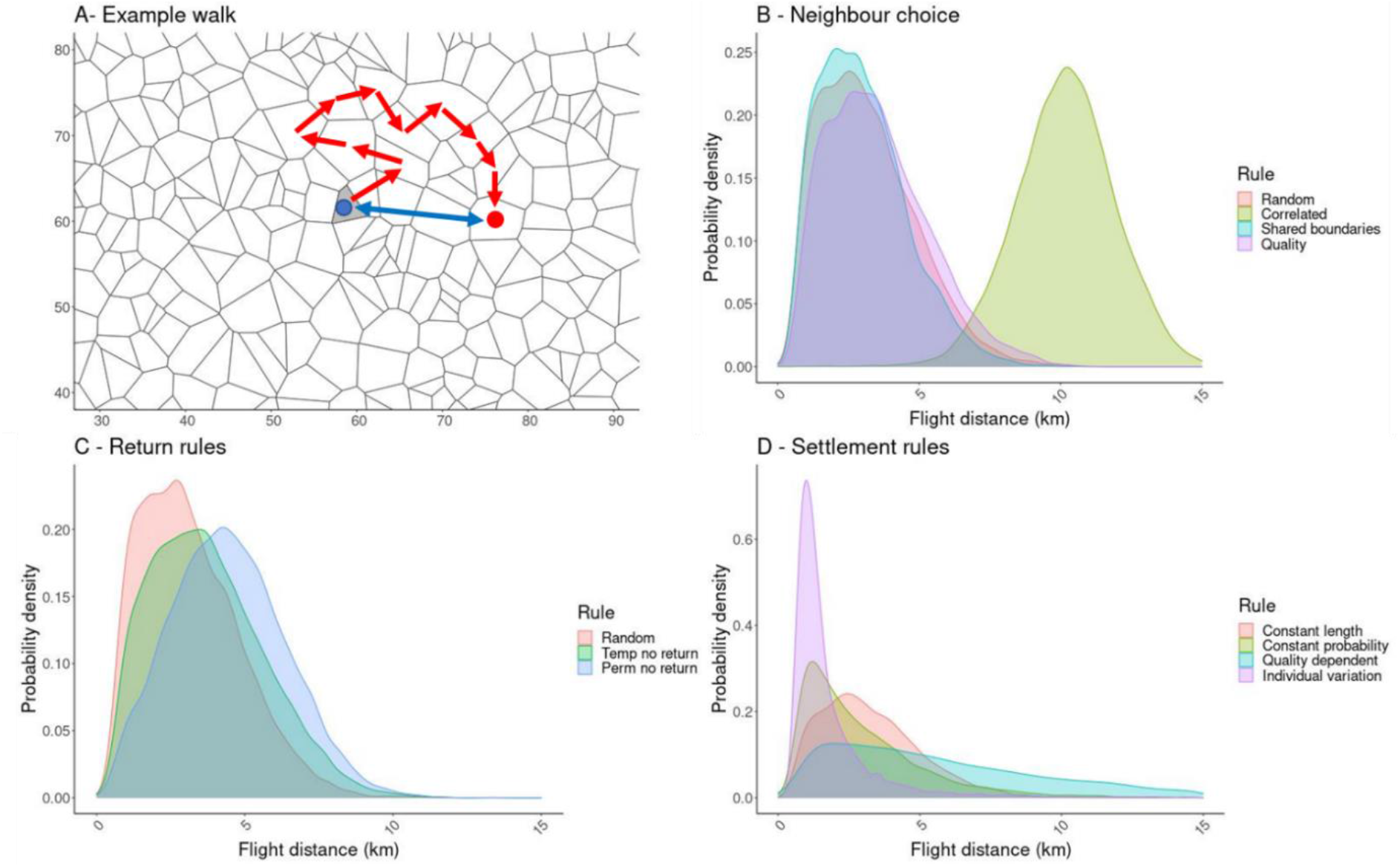
Influence of movement rule choices on the emergent dispersal kernel. Panel A shows an example of a 10 step walk through a mosaic landscape, red arrows indicate the walk and the blue arrow the flight distance. In each of the other 3 panels, the red kernel indicates our baseline rule set, approximating a discrete random walk (10 steps, no return rules and random neighbour choice). Relative to this, panel B shows the effect of neighbour choice rules, C return rules and D settlement probability rules.

For each rule combination, we characterised the resulting dispersal kernel using four criteria: mean flight distance (average Euclidian distance between centroids of origin and final settlement parcels); skew (dispersal kernels should be right-skewed as most individuals remain relatively close to their origin); kurtosis (how fat-tailed the kernel is, representing the probability of long-distance dispersers) and the ratio of mean walk length (sum of the centroid-centroid distances for each step) to mean flight distance (walk: flight). When calculating these measures, we excluded walks which ended in the original parcel as dispersal did not occur (therefore for some rule combinations, total number of walks <10,000). While kernels are expected to be highly variable depending on the specific species and landscape, typical kernels show high levels of variation (Nathan et al., 2012), being right skewed (Skewness>0) and heavy tailed (Kurtosis >3, more heavy-tailed than normal distribution). We therefore consider which combinations of rules generate these characteristics.

### Establishing fox kernel distributions for comparison

To demonstrate how movement rules may be selected for simulating dispersal, we take a realistic kernel, estimated from empirical data on red fox dispersal, and identify rule sets which approximate this kernel. Foxes have been studied extensively, being a widespread species and implicated in rabies transmission and maintenance. Fox capture-mark-recapture data shows that most foxes disperse only short distances (typically <10 km in urban areas (Harris & Trewhella, 1988)), but long-distance dispersal also occurs, and could be underestimated due to the limitations of dispersal studies (Nathan et al., 2008; Van Dyck & Baguette, 2005). Dispersal distances differ by sex, with males dispersing significantly further than females (Trewhella, Harris, & McAllister, 1988). Dispersal distance is also dependent on population density, with individuals at high population densities having smaller home ranges and dispersing shorter distances. We aimed to recreate the dispersal kernel for males in a high-density population typical for the UK. Parcels were generated typifying territory sizes observed in non-rural areas of the UK, of 1 km^2^ (Trewhella et al., 1988). In a review of dispersal distances for foxes, Trewhella (1988) found that mean dispersal distance for male foxes relative to home-range size can be estimated according to:

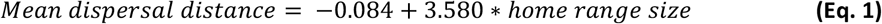

Both Trewhella et al. (1988) and Whitmee (2012) found that dispersal distances for foxes approximated a negative exponential distribution. Following Equation 1 we aimed to recreate an exponential distribution with λ=1/3.5km, based on a home range size of 1 km^2^.

In addition, to test the flexibility of the method, we also compared the exponential kernel based on empirical evidence against two alternative kernels of the same approximate scale but differing shapes (Table 2). Where there is a greater probability of long-distance dispersal, a more fat-tailed kernel is required, represented here with a half-Cauchy distribution with a modal value of 3.5 km (σ=3.5). Balancing this we also considered a distribution with a thinner tail than the exponential, using a gamma distribution with shape parameter of 5 and scale of 3.5/5. Exponential, gamma, and half-Cauchy distributions are all commonly used to model dispersal kernels (Fandos et al., 2021; Nathan et al., 2012).

**Table 2:**
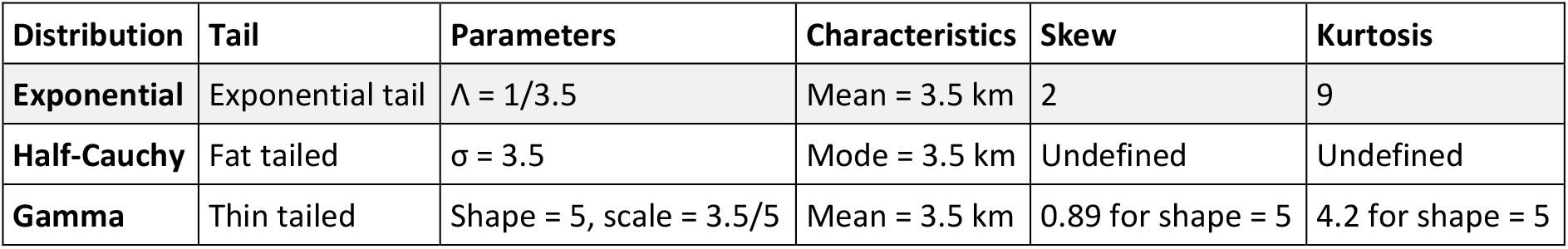
Example distributions for dispersal kernel. The distance distributions produced were compared to a kernel based on empirical evidence (Exponential-highlighted in grey) and two alternative distributions (Half-Cauchy and Gamma).

| Distribution | Tail | Parameters | Characteristics | Skew | Kurtosis |
| --- | --- | --- | --- | --- | --- |
| **Exponential** | Exponential tail | $\Lambda = 1/3.5$ | Mean = 3.5 km | 2 | 9 |
| **Half-Cauchy** | Fat tailed | $\sigma = 3.5$ | Mode = 3.5 km | Undefined | Undefined |
| **Gamma** | Thin tailed | Shape = 5, scale = 3.5/5 | Mean = 3.5 km | 0.89 for shape = 5 | 4.2 for shape = 5 |

The resulting distributions of flight distance for each combination were compared to the three different kernels using a Kolmogorov-Smirnov test, a non-parametric test of whether a sample comes from a particular distribution. For each distribution, the three best fitting emergent kernels (defined as having the lowest value for the Kolmogorov-Smirnov test statistic) were extracted and used to assess potentially suitable movement rule sets.

## Results

We assessed the kernels generated by dispersal movements within synthetic landscapes formed by Voronoi tessellation. This method of landscape generation produces parcels with a distribution of sizes, step-lengths, and neighbourhoods (see S1 Figure 1), as opposed to the use of regular grids in which these traits are fixed. Conceptually, parcels approximate to circles with mean area, mean centroid-to-centroid step lengths and mean neighbourhoods all conforming to a tessellation of circles. Consequently, the relationship between mean parcel area and mean step length (the average distance from centroid to centroid for steps of the dispersal walk) is equal to:

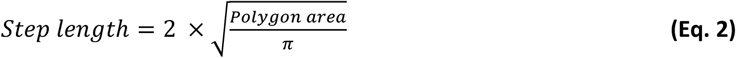

Therefore, for the simulated landscapes with a mean parcel size of 1 km^2^ (S1 Figure 1A), the mean step length will approximate 1.12 km (empirically validated in S1 Figure 1B). On average, each parcel has 6 directly adjacent neighbours (Range 1-11). Emergent distance kernels resulting from stepwise dispersal across these landscapes are a product of the step-length distribution (determined by the specific landscape), the distribution of number of steps taken by agents (determined by the settlement rules) and the distribution of angles of movement (determined by the neighbour choice and return rules). Relationships between these variables in mosaic-based landscapes are independent of scale. As a result, sets of dispersal movement rules applied across mosaics will produce consistent kernel shapes, albeit transformed with differing scales e.g., mean distances (See S1 Section 4 in the supplementary material for illustration). The dispersal rules considered here are also all applicable to regular geometries (e.g., grids). However, as step lengths and number of neighbours are fixed in these cases, the resulting kernel would differ, with less variation possible.

### Influence of dispersal rules

As expected, different kernels are generated depending on the dispersal rule set simulated. The baseline dispersal rule combination considered was a discrete version of a random walk for a set number of steps (random neighbour choice, no rules on return, 10 steps), producing a right-skewed kernel (mean = 3.2 km; skew=0.75) with a kurtosis of 3.3 (red kernels in Figure 1), close to that expected for a normal distribution. Relative to this baseline, of the neighbour-choice rules the correlated walk was most divergent (Figure 2B) as movement routes are straighter, leading to a substantially higher mean flight distance (10.1 km) and a less skewed distribution for the same number of steps (Skew=-0.08). By comparison, weighting neighbour choice based on local geographic factors (length of shared boundaries or parcel quality) produced a kernel broadly comparable to a random walk in this study, although the latter will be highly dependent on how landscape heterogeneity (e.g., resistance to movement) is represented and the character of the landscape (e.g., well connected, fragmented).

Prohibiting agents from selecting previously visited parcels extended mean dispersal flights compared to our random walk (Figure 2C). In our limited exploration of this behaviour, a reduced tendency to revisit parcels produced greater mean flight distances for the 10-step fixed-length walks (mean flight distance 3.2 km for temporary no return, relative to 4.5 km for persistent no return). Furthermore, the walk to flight ratio changed progressively (4.9 vs 4.1 vs 3.2) as less tendency to revisit parcels in general reduced the tortuosity of walks. The shape of kernels also changed progressively with reducing skew as the tendency to return decreased (Skew: no return rules 0.74, temporary no return 0.61, persistent no return 0.31).

For settlement rules, including stochasticity in the number of steps taken by using a settlement probability function produced greater skew in the emergent dispersal kernels (Figure 2D: Constant walk skew 0.8 vs fixed settlement 1.7) and dramatically different maximum flights (Constant walk 10.9 km vs fixed settlement probability 20.3 km). The effect of allowing quality-dependent settling, rather than assuming a homogenous landscape, will be dependent on the specific landscape simulated. In this case, settlement dependent on quality led to a comparable skew and kurtosis to a constant settlement probability (skewness= 1.8 and kurtosis =7.5), but mean distance was substantially higher at 6.6 km. Increasing variability further by allowing individuals to vary in their settlement probability (In this case, sampled from a uniform distribution between 0 and 1) had a substantial impact on the resulting kernel. For individual variability, the skew (6.1) and kurtosis (63.0) of the resulting kernel were greatly increased, resulting in a maximum dispersal distance of 43.3 km.

### Sensitivity analysis

Interactions between different rules and their specific parameterisations may lead to different walk behaviour, and therefore resultant kernels. A full comparison of kernel characteristics between all 40 rule sets across a range of parameterisations is presented in S1 Figure 3-5. Here we restrict ourselves to a qualitative description of the main effects and interactions. Our aim was to identify rule sets that may provide flexibility for modellers aiming to recreate plausible dispersal kernels.

**Figure 3:**
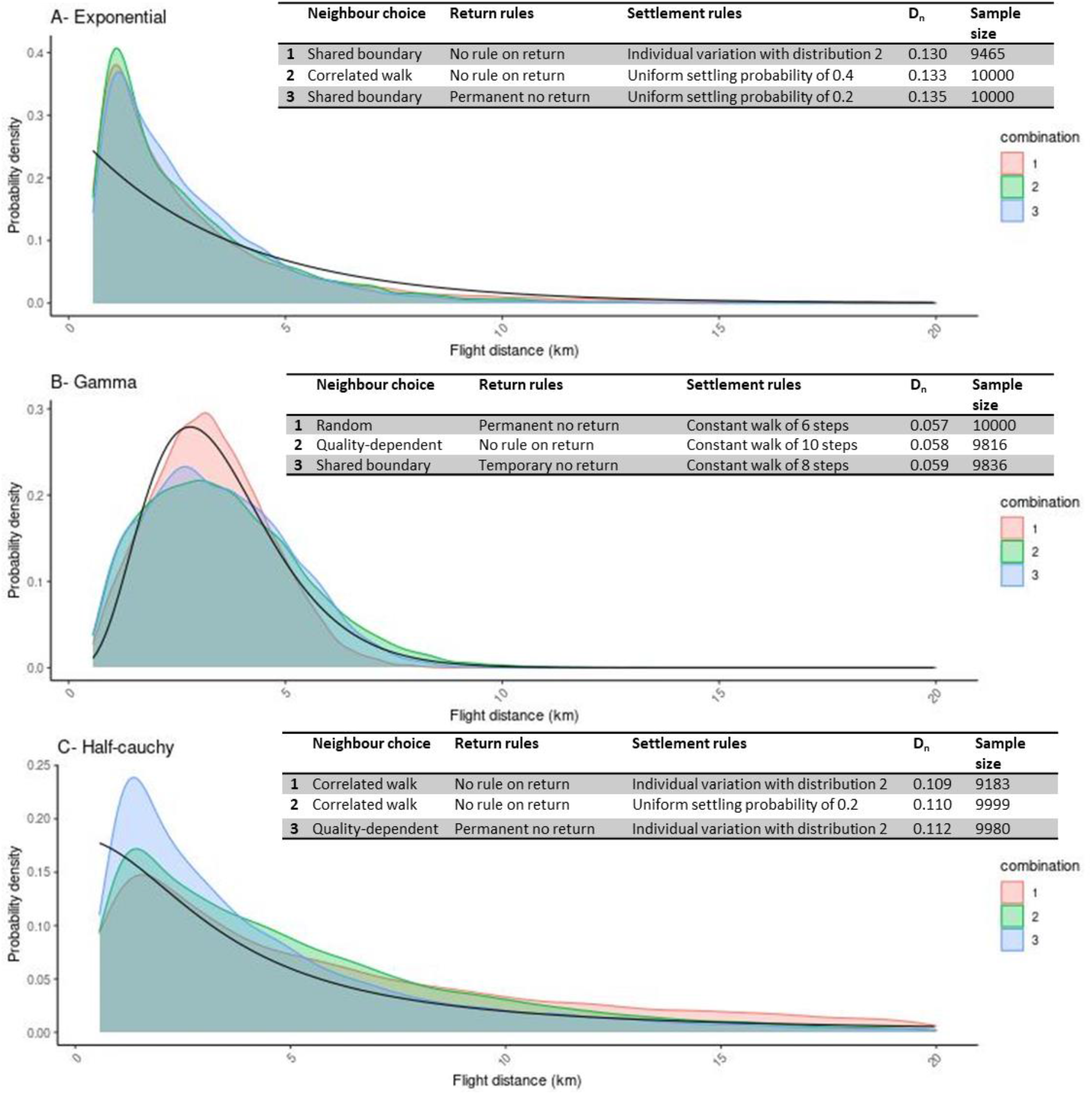
Best fitting rule sets to reference distributions representing red fox dispersal. The reference kernel is shown in black, with probability values shown from a range of 0.56 (mean parcel circumference-minimum distance considered to have dispersed) to 20 km. Colours indicate which movement rule set was used, which is presented in the tables.

Results of the sensitivity analysis highlighted the substantial influence of settlement rules on the dispersal kernel, relative to neighbour choice and return rules. In general, neighbour choice did not have a substantial influence on the resultant kernels, with the exception of rule sets including correlation substantially increasing the mean dispersal distance (S1 Figure 3). Incorporating return rules also had a consistent effect when combined with any other rules, of a relatively marginal increase in mean dispersal distances. While return rules and neighbour choice rules influence the movement route taken, they have no impact on the number of steps taken, only the walk to flight ratio, and therefore have limited influence on flight distances.

To recreate a skewed distance distribution with a heavy tail (which are correlated traits as shown in S1 Figure 6) the choice of settlement rules applied to agents and their parameterisation is highly influential as this determines the variation in the number of steps taken across the entire walk. The simplest rule considered, of a set number of steps, can only generate kernels with low skew and kurtosis, regardless of which return and neighbour choice rules are used (S1 Figures 4A and 5A). Introducing stochasticity by either incorporating heterogeneity in parcel descriptions and allowing agents to respond to these (here quality-dependent settlement probability, S1 Figures 4C and 5C), or by allowing consistent individual variation in their dispersal walks (S1 Figures 4D and 5D), allowed for much greater variation in the resultant kernels. Of the rules considered, incorporating individual variation led to the most highly skewed kernels. In particular, this was greatest in populations of agents where a substantial proportion had a low per-step probability of settlement (e.g., due to adventurous personalities or high energy reserves allowing longer movements before settlement). However, it should be noted that we only considered limited variation in parcel quality and changing the quality distribution and response to this would further influence the resultant kernel.

### Movement rules for reference fox kernels

The empirical distribution of distances travelled by foxes during dispersal can be represented by an exponential distribution. Comparison of movement rules found that this distribution was best recreated (defined by the lowest test statistic for the Kolmogorov-Smirnov test) by a movement rule allowing individual variation in settlement probabilities (with each agent’s settlement probability randomly chosen from a beta distribution with α=1, β=5), with neighbours selected based on the length of shared boundaries and no restrictions on returning to previously visited parcels. However, alternative rule sets also provided similar fits to this distribution, despite making substantially different assumptions about movement behaviour, for example, a correlated walk with a constant settlement probability (Figure 3A) also provided a good fit to the empirical distribution. In no cases were perfect fits achieved, and for all rules, p-values were less than 0.001, showing distributions were significantly different from the reference, as may be expected for the large sample sizes used (>9400 individual flight distances in all cases). However, from visual inspection of Figure 3, it can be seen that acceptable recreations were achieved, allowing movement rules to capture key features of fox dispersal.

For the half-Cauchy distribution, the best fitting movement rules were provided by either allowing individual variation or a uniform settlement probability, which provide the fat-tail characteristic of this distribution. By contrast, for the gamma distribution, settlement after a constant number of steps was selected as this led to less skew in the resulting distribution. As with the exponential distribution, while perfect fits were not achieved, the movement rules were able to capture the general shape and scale of these reference distributions, demonstrating the flexibility of the rules considered to generate a range of kernel shapes, despite their simplicity. Notably, while the best-fitting rule sets for each kernel provided very similar flight distance distributions, the distribution of walk distances differed substantially (S1 Figure 8), as these walks differ in their walk to flight ratio.

## Discussion

Spatially explicit mechanistic models are a useful tool to inform decision making through prediction of previously unobserved or unquantified events. In the context of wildlife, areas of interest for predicting spatial dynamics include native species management (and the deployment of novel interventions), spread of invasive species (especially those known to cause harm and where immediate eradication is mandated), and the costs and mitigation of notifiable disease (White et al., 2018b). Models for these purposes generally involve a representation of a landscape, within which animals follow mechanistic behaviours, such as reacting to one another and their local environments (collective processes representing their population dynamics), and across which they move. The development of such models to inform decision-making requires modelling concepts to be flexible and robust when projected into real-world geographies and used to represent real species ecology. In the absence of directly applicable validation data, demonstrating their utility depends on the assurance that model abstractions and processes are fit-for purpose and do not produce bias. This study considers the suitability of methods for modelling individual dispersal movements across an irregular mosaic landscape (a type of geographic automata), in order to represent dispersal dynamics of terrestrial vertebrates. We explore the behaviour of a set of simple rules governing the movement of an agent and show how the resulting dispersal kernels may be used to corroborate the rule sets chosen. The most appropriate rule set will be highly specific to the particular species and landscape; therefore, our aim was not to prescribe a particular method but to illustrate a range of basic alternatives, demonstrate qualitative features of the emergent kernels and how these may be compared to a real-world example.

A broad range of dispersal kernels emerged from the rule sets considered, with differences which could have substantial consequences for model behaviour. Notably, the most basic rule type considered, an approximation to a random walk, produced a kernel showing little skew and close to a normal distribution. While the simplicity of this rule may be attractive in some modelling contexts, our results highlight the limitations for use in recreating a “typical” mammalian dispersal kernel, which tend to be highly leptokurtic (Nathan et al., 2012). The more complex movement rules considered made different approximations to the real behaviour of animals (See Table 1). Inclusion of these behavioural approximations can generate kernel shapes closer to the “typical” distance distribution. Hawkes et al. (2009) reviewed factors that may contribute to the skew observed in empirical dispersal kernels, including population heterogeneity, landscape features and correlation. While our results cannot inform the causative factors behind leptokurtic distributions, in agreement with Hawkes et al. (2009), of the rules we consider, those including heterogeneity in settlement probability due to individual variation or landscape heterogeneity most effectively generated skewed and heavy-tailed kernels. Using the example of red fox dispersal, the best fitting movement rule assumed individual variation in settlement, with agents able to return to any previously visited parcel and selection of neighbours based on shared boundary length. While a perfect fit to the reference kernel was not obtained, the resulting distribution showed that even simple assumptions about movement behaviour can recreate a plausible kernel and would produce acceptable simulations.

As dispersal methods seek to emulate real species-specific behaviour, where prior information on dispersal behaviour is available this should be incorporated. Using our fox example, for the exponential kernel, a correlated walk with a constant settlement probability provided almost an identical fit compared to the best-fitting movement rule set, despite making substantially different behavioural assumptions. This finding emphasises the need to use further insight into the movement biology of study species where available. In foxes, dispersal events tend to be straighter than other movements as this is thought to improve dispersal success (Soulsbury et al., 2011; Zollner & Lima, 1999). Therefore, a correlated walk may be more realistic than neighbour choice based on shared boundaries. However, in many cases there may be little information on disperser behaviour and its drivers. In these cases, choosing a set of simple rules that generates plausible kernels may be more appropriate than rules requiring additional assumptions without appropriate data, especially where those simple rules may also reduce the potential for bias and artefact. Evidently, we do not consider the full range of dispersal rules that could be applied in a spatial model. There are countless potential combinations and parameterisations that could be used to capture the subtleties of animal movement behaviour. Although consideration of additional rules could lead to improved fits to empirical kernels, our aim was not to be exhaustive but to highlight potential options and how these can influence the resulting dispersal walks.

The emergent kernel from dispersal simulations is a product of both the landscape and the dispersal rule set applied across it. In our framework, the landscape supplies a distribution of step-lengths, the neighbourhood, and parcel qualities, whilst the rule set defines the number of steps taken and the direction of each, producing variation in the tortuosity of dispersal walks and the walk to flight ratio. In our irregular mosaic, these factors are scale-independent, as increasing the parcel size for a randomly generated mosaic will influence the magnitude, but not the distribution, of step lengths, and consequently the shape of kernels is scale-free. For some species (as in our fox example), parcel size may be determined by known variables such as territory size (Trewhella et al., 1988). However, where there is a decision to be made on appropriate parcel size, if the set of movement rules is pre-established and there is an estimate of the scale of dispersal distance (e.g., mean distance) it is possible to back-calculate an appropriate mean parcel size to produce the necessary kernel scale (See S1 Section 4 in the supplementary material). Here we note that parcel sizes that are too large (coarse grain) may provide insufficient steps between an origin and the mean dispersal distance to generate sufficient variation in the resultant kernel, whilst parcels that are too small (many steps between origin and mean) may be unnecessarily computationally demanding. As such, the scale of dispersal movements relative to the parcel size should be one consideration for modellers during the design of the spatial representation, in addition to other factors such as the scale across which daily movements and resource acquisition occur.

In this exploratory study, we used a simplified synthetic landscape consisting of randomly generated parcels. By generating parcels independently of the underlying habitat quality, we assumed these characteristics were unrelated. However, for species where territory sizes are known to depend on habitat quality and resource availability, such as the red fox (Trewhella et al., 1988), this might be incorporated when generating the model landscape. Including quality-dependent parcel sizes would influence the local step-length distribution and therefore the resulting kernel. In our fox example, this assumption can capture the observation that dispersal distances are much shorter in urban areas where territories are small, and they are densely packed due to higher resource availability (Trewhella et al., 1988). The link between parcel size and dispersal distance allows this relationship to be recreated without additional assumptions about fox behaviour in rural or urban areas. A further simplifying assumption used in this paper was the absence of barriers restricting dispersal movements. However, our method of landscape generation, using irregular geometry, allows for realistic representation of barriers, regardless of their scale (Holland et al., 2007). The influence of barriers will be highly specific to the particular landscape. For this reason, we would always advocate that processes running across landscapes are verified to ensure they do not produce bias when used ‘*in situ*’ (representations of real landscapes). However, by exploring their properties in a simple synthetic landscape with known properties we establish that the movement process is not fundamentally prone to artefact and is verified as fit for purpose.

In this paper, we have demonstrated the variable kernels generated by movement rules and how they can be corroborated against empirical information. Corroborating movement rules is important given the potential that dispersal has population-level impacts on model outputs. For example, movement routes may have substantial impacts on dispersal-mediated disease spread, with longer and more torturous walks leading to more potential contacts. Responses of dispersers to landscape features may also be of substantial influence. For example, Scherer et al. (2020) showed that compared to a correlated walk, movement based on response to landscape heterogeneity increased pathogen persistence, as the resulting spatial heterogeneity in infection probabilities (e.g., hotspots of infection in high quality areas) maintained sub-populations of both susceptible and infectious individuals. For this reason, where available, detailed information on species movements and preferences can also be used to inform the “quality” layers and improve model predictions. In this paper, we considered a single abstract quality which varied across the landscape. In reality, animals are likely to trade-off decisions using a range of ‘quality’ factors such as resistance to movement, habitat suitability, resource availability, reproductive competition, and predation risk. Modelling studies are increasingly incorporating detailed landscape information influencing species-specific movement (e.g., Graf, Kramer-Schadt, Fernández, & Grimm, 2007; Kanagaraj et al., 2013). Movement rules may also influence population-level processes when incorporating disperser mortality. If dispersers face a mortality risk at each step, longer dispersal events are more likely to result in death (rather than successful settlement and breeding), relative to shorter, more directed movements, with all else being equal. Dispersal mortality is therefore an additional potential source of variation in kernels with impacts on population survival (Hein, Pfenning, Hovestadt, & Poethke, 2004). Overall, dispersal is a complex process influenced by numerous factors. Identifying these drivers for particular species is highly challenging, requiring intensive empirical studies. However, by verifying movement rules against available information, such as dispersal kernels, the reliability of predictive mechanistic processes and the models they inform may be improved, even where the full underlying process is not understood.

## Conclusions

Realistic representation of dispersal is central to predicting spatial dynamics of animal movement across landscapes. We demonstrate the ability to represent dispersal by simple movement rules across an irregular mosaic landscape. Our method of landscape representation is free of bias in movement directions and can be adapted to allow accurate representation of barriers and underlying landscape patterns. When simulating movement across these landscapes, a broad range of options for movement rules are available to the modeller. Our results show that particular rules, such as allowing variation in settlement dependent on underlying landscape quality or individual-level variation, may allow more realistic movement patterns. However, rules will be highly specific to the species and particular landscape and should therefore be validated within the specific modelling context used.

## Supporting information

S1 File - Additional Outputs

## Acknowledgment

This work was funded by the Department for Environment, Food and Rural Affairs (Defra) under project code EXSE0432.

