## Supplementary material for "Selection of movement rules to simulate species dispersal in a mosaic landscape model": S1 File - Additional Outputs

### Appendix 2: Additional model outputs

#### 1. Parcel characteristics

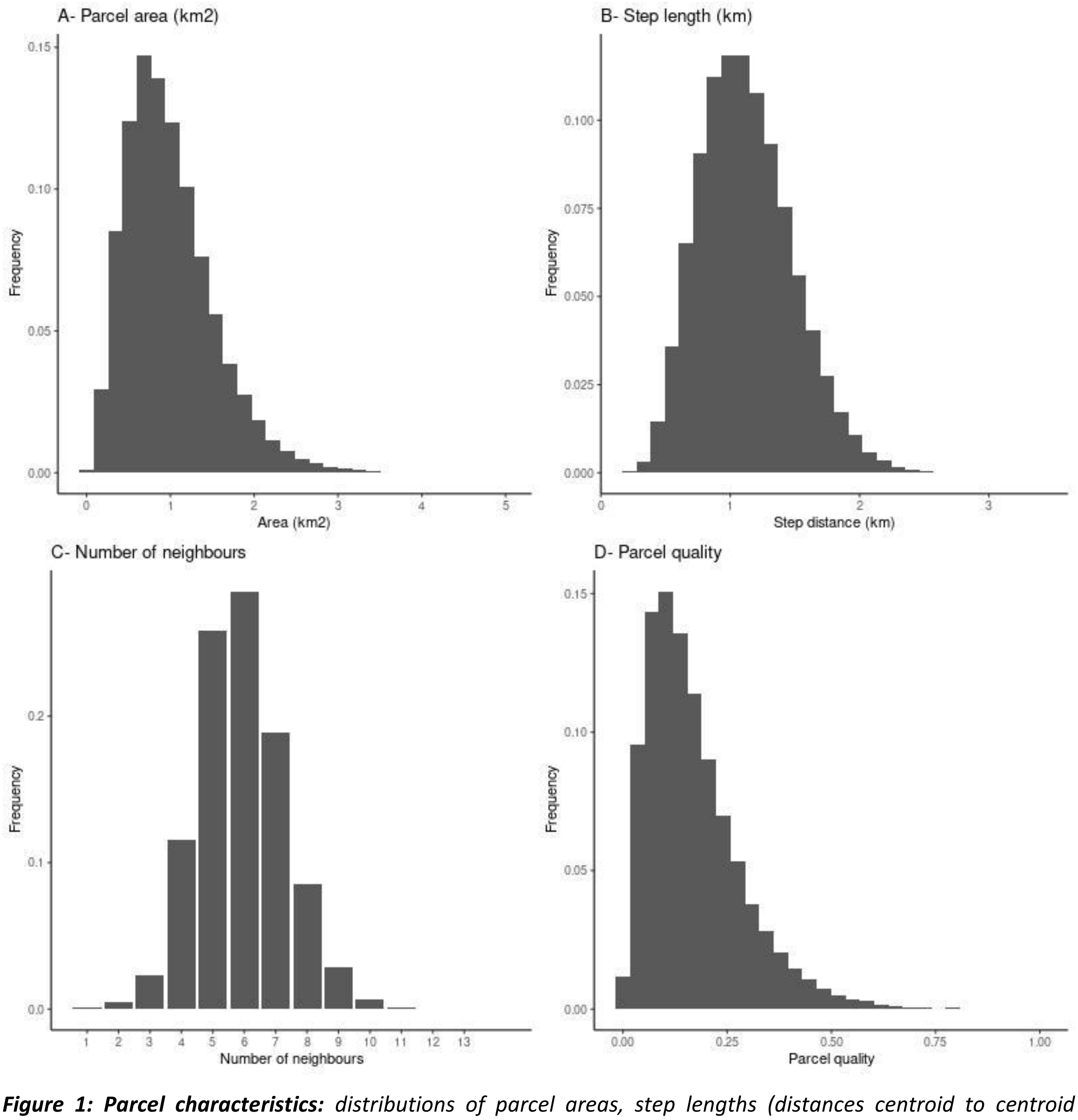


***Figure 1: Parcel characteristics:*** *distributions of parcel areas, step lengths (distances centroid to centroid between neighbours), number of neighbours and relative parcel quality are shown.*

#### 2. Distributions for individual variation in settlement probability

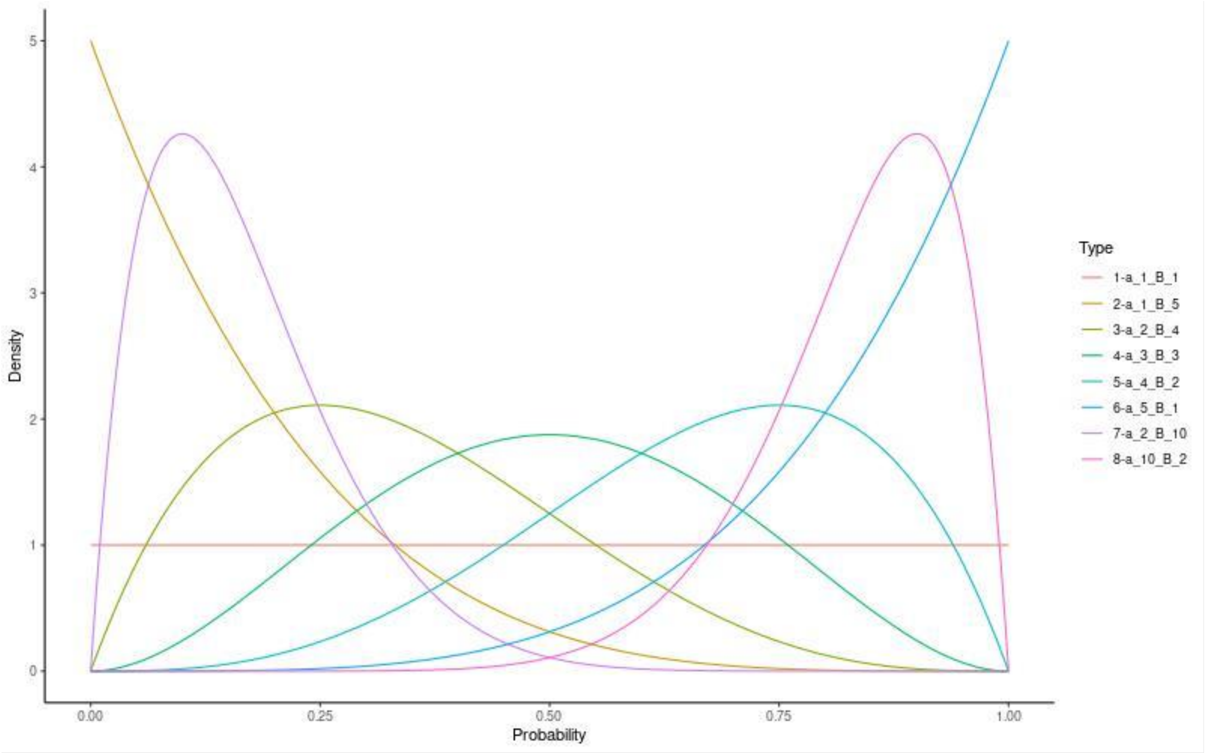


***Figure 2: Potential beta distributions for probability of individual settlement per step.*** *In this context, a right-skewed distribution produces lower probabilities of settling, leading to longer dispersal walks on average.*

#### 3. Sensitivity analysis results

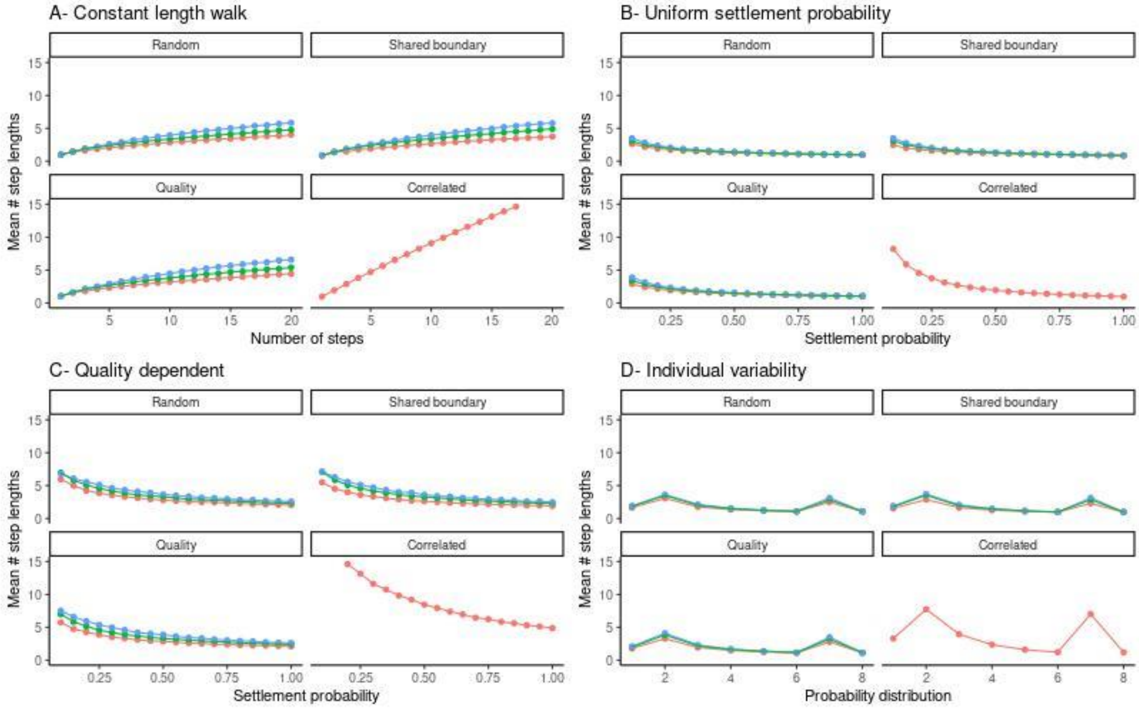


***Figure 3: Influence of walk type and input parameter on mean distance travelled relative to mean step length.*** *Colour indicates return rules: green=no return rules, blue=temporary no return, red=permanent no return.*

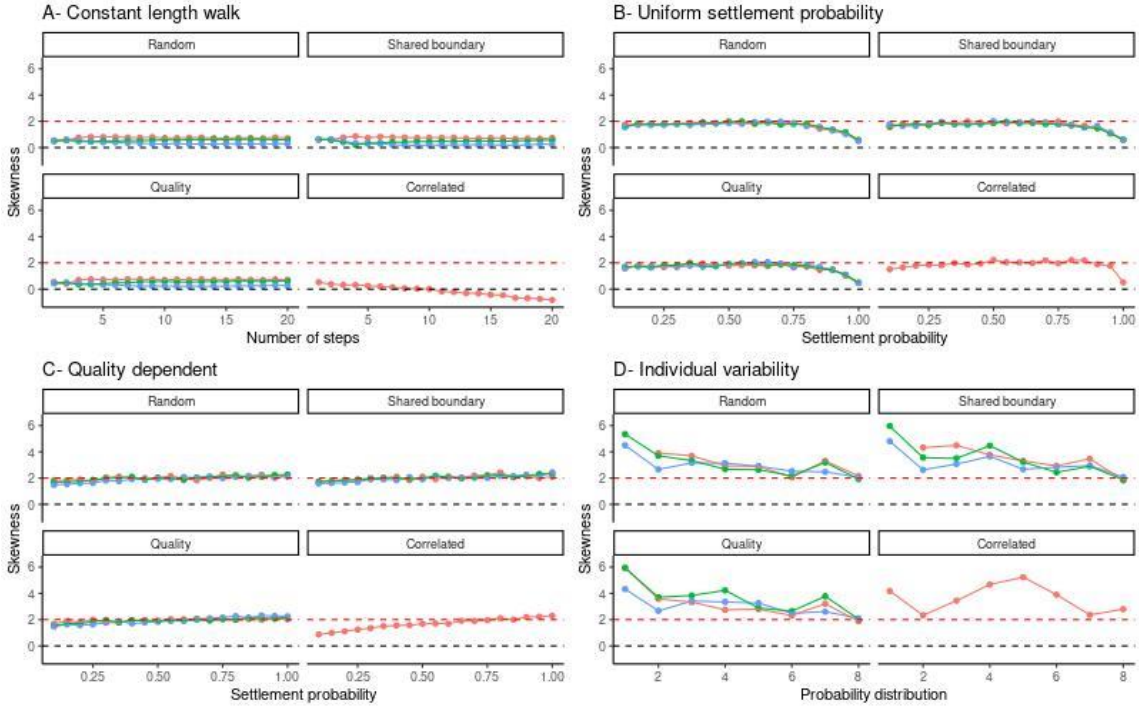


***Figure 4: Influence of walk type and input parameter on skewness of resulting dispersal kernel.*** *The black dotted line indicates a skewness of 0, equivalent to a symmetrical normal distribution, and the red dotted line a skewness equal to 2, equivalent to an exponential distribution. Colour indicates return rules: green=no return rules, blue=temporary no return, red=permanent no return.*

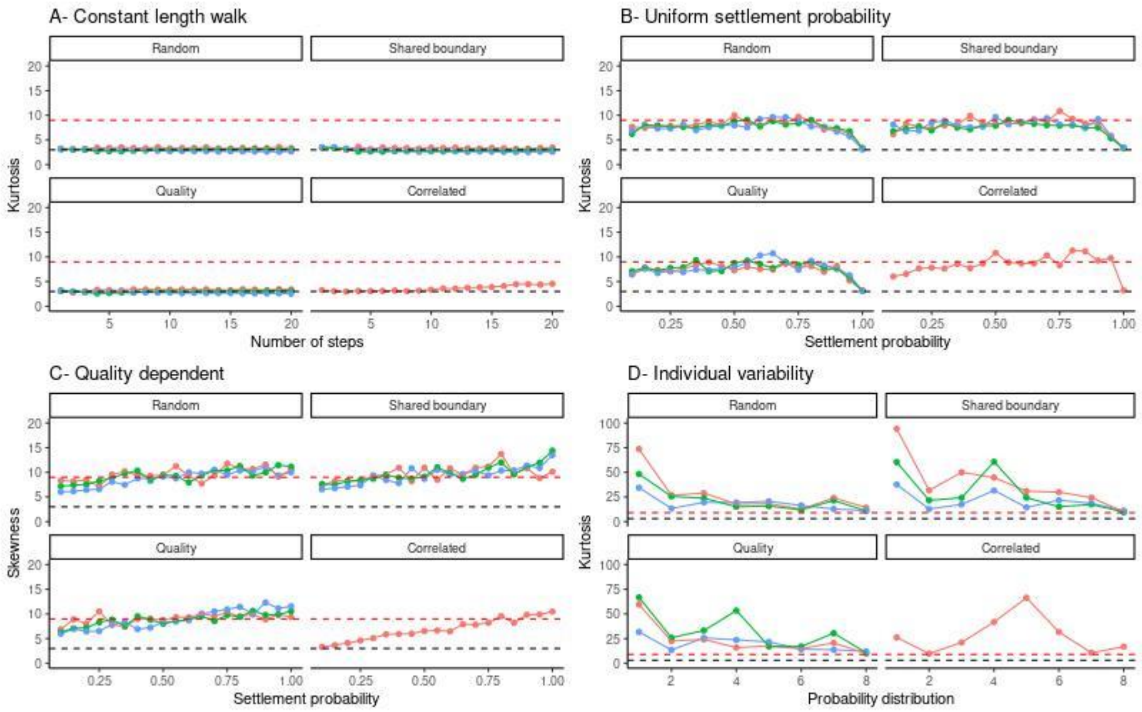


***Figure 5: Influence of walk type and input parameter on kurtosis of resulting dispersal kernel.*** *The black dotted line indicates a skewness of 3, equivalent to a normal distribution, and the red dotted line equal to 9, equivalent to an exponential distribution. Note the scale is between 0-25 for Fig. 5A-C and 0-100 for D. Colour indicates return rules: green=no return rules, blue=temporary no return, red=permanent no return.*

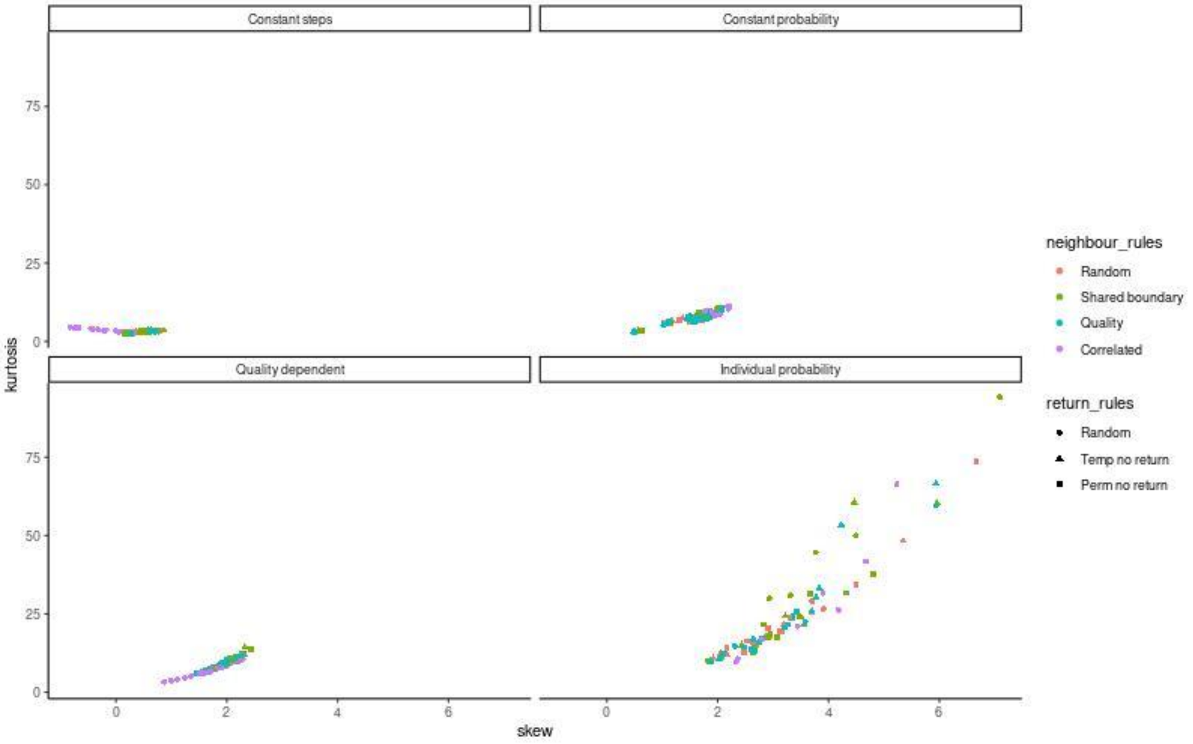


***Figure 6: Relationship between skew and kurtosis for all possible walk combinations.*** *Each point represents a particular walk combination with a single parameterisation.*

#### 4. Back-calculating parcel size for a particular movement set

In the main text, we consider movement rules for red fox, based on selecting a best-fitting kernel within a landscape of a chosen parcel size pre-determined by an assumed mean fox home range size. If parcel size is not pre-specified, it is also possible to choose an initial set of movement rules and adjust the scale of parcel sizes to fit to the target kernel.

As an example, we assume a pre-specified rule set for fox movement: a correlated walk of constant length (10 steps), with no rules on return. As shown in Figure 3, this walk generates a mean flight distance of 9.15 times the step length. For a landscape with a pre-specified mean parcel size of 1 $km^2$ and step length of 1.12 km, this therefore generates a mean flight distance of 10.25 km, substantially further than the empirical estimate (3.5 km). To fit the scale of the dispersal kernel to the empirical estimate, the scale of the parcels can be adjusted.

To give a mean flight distance of 3.5 km (the empirically estimated mean), for this particular rule set a step length of 0.38 km (3.5/9.15) is required, equivalent to a mean parcel size of 0.11$km^2$ (based on polygons approximating circles when averaged). To generate a landscape of 10,000 parcels with this specified size, a total arena of 1,148 $km^2$ (33.9 km by 33.9 km) is required. Running the same rule set in this re-scaled landscape produces the same shape kernel as in the original landscape (Fig. 7B) but is scaled so that the mean dispersal distance is 3.5 km.

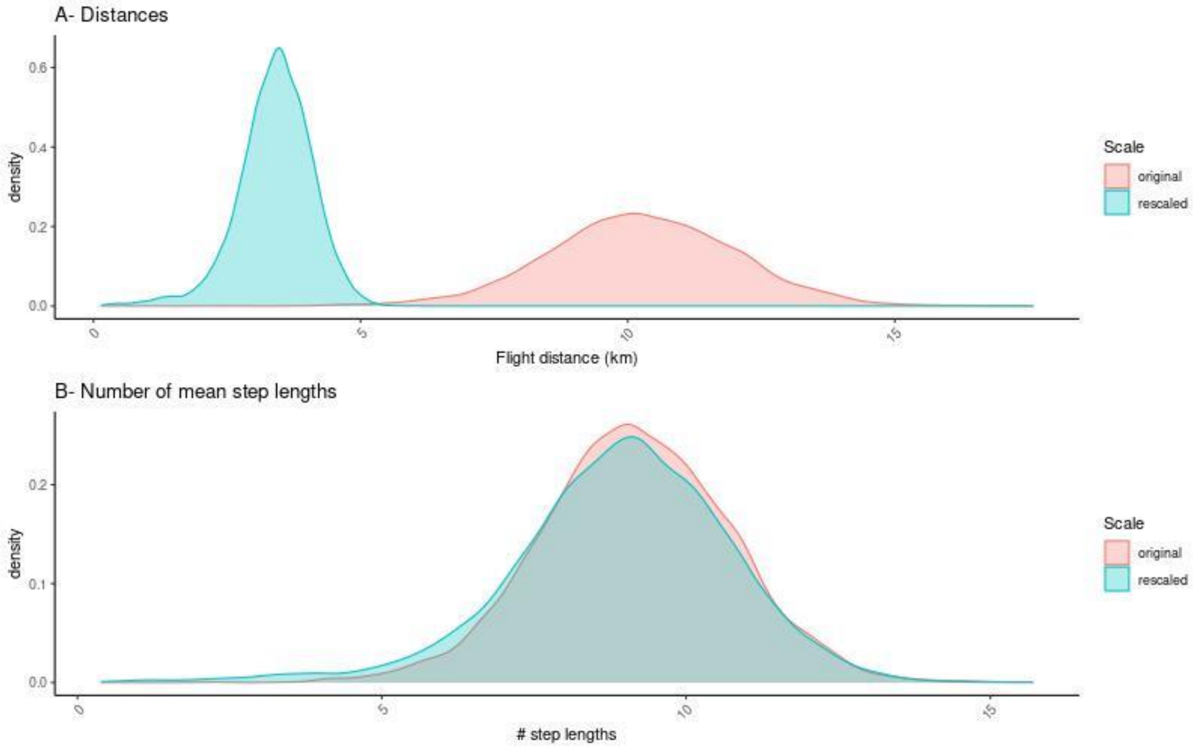


***Figure 7: Example of rescaling landscape to adjust scale of resultant kernel.*** *The same rule set (correlated walk of 10 steps with no rules on return) was run in a landscape of a mean parcel area of 1 $km^2$ (original) and a landscape with a mean parcel area of 0.11 $km^2$ (rescaled). Relative to the step length, the same kernel is produced (B) but the scale (actual distance) differs (A).*

#### 5. Walk distances for fox reference kernels

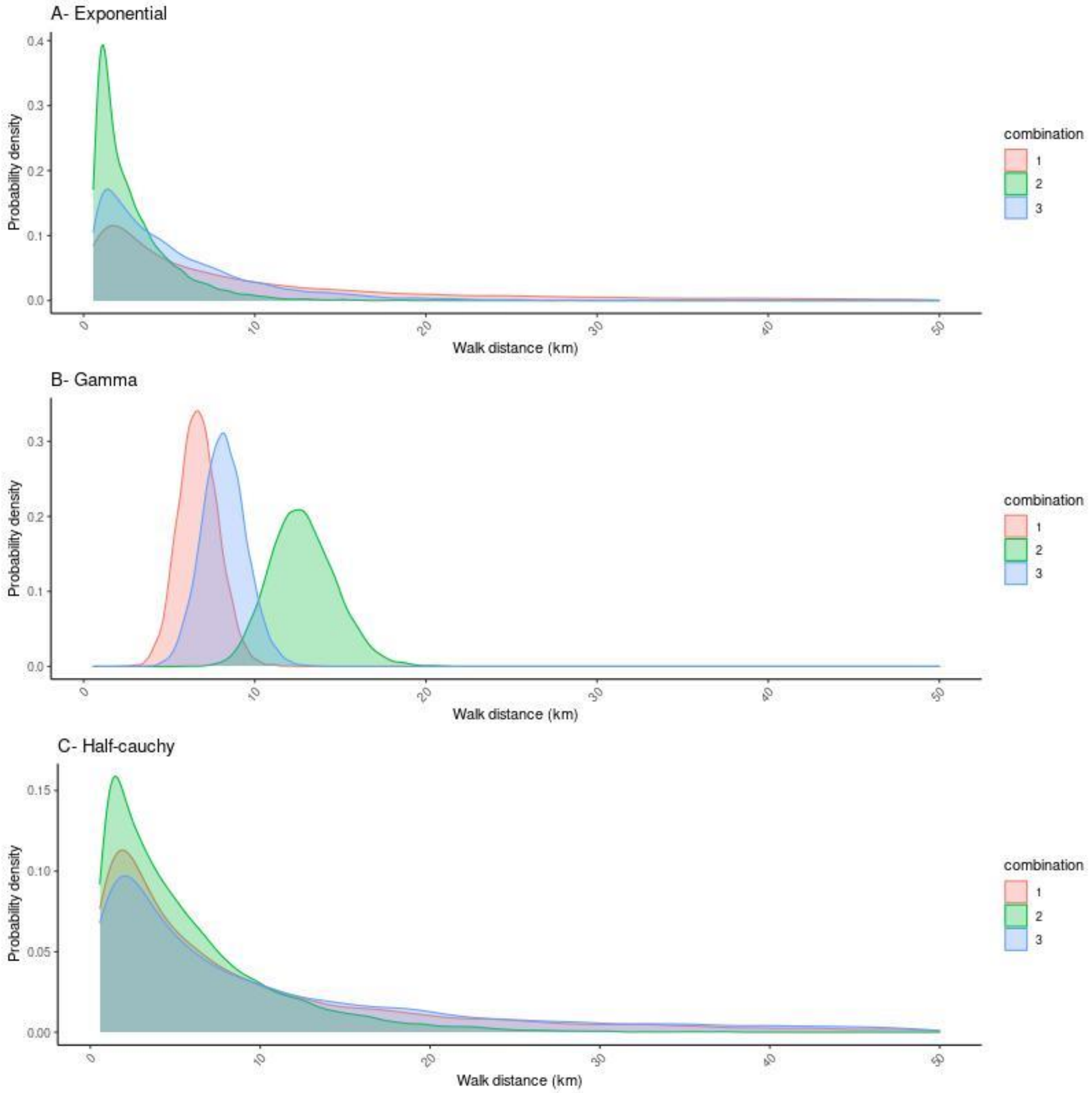


***Figure 8: Walk length distributions (cumulative length of parcel-to-parcel movements) for each of the best fitting rule sets for fox reference kernels shown in Figure 3 in the main text.***
